# Karyotype-environment associations support a role for a chromosomal inversion in local adaptation in the seaweed fly *Coelopa frigida*

**DOI:** 10.1101/278317

**Authors:** Claire Mérot, Emma Berdan, Charles Babin, Eric Normandeau, Maren Wellenreuther, Louis Bernatchez

## Abstract

Large chromosomal rearrangements are thought to facilitate adaptation to heterogeneous environments by limiting genomic recombination. Indeed, inversions have been implicated in adaptation along environmental clines and in ecotype specialisation. Here, we combine classical ecological studies and population genetics to investigate an inversion polymorphism previously documented in Europe among natural populations of the seaweed fly *Coelopa frigida* along a latitudinal cline in North America. We test if the inversion is present in North America and polymorphic, assess which environmental conditions modulate the inversion karyotype frequencies, and document the relationship between inversion karyotype and adult size. We sampled nearly 2,000 flies from 20 populations along several environmental gradients to quantify associations of inversion frequencies to heterogeneous environmental variables. Genotyping and phenotyping showed a widespread and conserved inversion polymorphism between Europe and America. Variation in inversion frequency was significantly associated with environmental factors, with parallel patterns between continents, indicating that the inversion may play a role in local adaptation. The three karyotypes of the inversion are differently favoured across micro-habitats and represent life-history strategies likely maintained by the collective action of several mechanisms of balancing selection. Our study adds to the mounting evidence that inversions are facilitators of adaptation and enhance within-species diversity.

## Introduction

Adaptation to heterogeneous environments is a major driver of evolution and in the diversification of life [1,2]. When a species occurs over a large geographic range, it experiences spatially variable conditions. Under limited migration, each population may thus follow its own evolutionary trajectory driven by local environmental conditions [1,3,4]. This can result in genetic and phenotypic polymorphism, variation at adaptive traits between habitats or along environmental clines, and ultimately, diversification into ecotypes or even species [3,5–8].

The ability to undergo polygenic specialisation and local adaptation is related to the amount of genetic exchange between populations, because gene flow mixes unfavourable immigrant alleles with resident alleles. In this context, negative modifiers of recombination can play an important role by limiting allele shuffling in parts of the genome [9–12]. Chromosomal inversions modify recombination because the heterozygote gene order is reversed between the standard and the inverted arrangements, resulting in limited recombination within the inverted part [13,14]. Various models have argued that inversions can facilitate local adaptation when they trap a set of co-adaptive alleles [15–18], conditions which are not unusual when inversions span hundreds of genes [19].

Empirical evidence supporting a link between inversions and local adaptation comes from clines of inversion frequencies along environmental gradients [16,20,21] or chromosomal rearrangements associated with ecotype divergence [22–25]. The pattern of inversion frequency distributions can be variable, ranging from fixation between distinct habitats [24,26], clinal modulation of intermediate frequencies [20] to widespread polymorphism in other cases [27–29]. The fate of an inversion depends on the selective mechanisms at play and on the genes trapped within the inversion. Strong selection and steep differences between habitats can drive inversions near fixation when locally-adaptive alleles are involved [17] and lower heterokaryotype fitness may drive divergence when populations carry alternative rearrangements [24,26,30]. By contrast, polymorphisms may be maintained by gene flow or if balancing selection is involved [29–33]. To disentangle these mechanisms, it is thus useful to combine knowledge of inversion effects on phenotypes with ecological and molecular data on natural populations harbouring the inversion across environmental gradients.

The seaweed fly *Coelopa frigida* provides a suitable model to understand how inversions facilitate adaptation to heterogeneous environments because it carries a large inversion on chromosome I, with the two forms called α and β [34], that is polymorphic in all populations sampled so far. Studies on European *C. frigida* have shown that the inversion frequency varies along a latitudinal cline in Scandinavia which follows a natural gradient of temperature, salinity and seaweed composition, that the inversion has large phenotypic effects, particularly on male size, development time, viability and fertility, and that heterokaryotes generally have a higher egg-to-adult viability than homokaryotes [35–40].

Here we investigate American populations of *C. frigida* along the North Atlantic coast. The sampled area follows a latitudinal cline, a gradient of salinity into the St Lawrence Estuary and spans an heterogeneous seaweed distribution. This allowed investigating separately the effects of different environmental parameters on the inversion frequency and testing the extent of parallelism with Europe. Specifically, we set out to determine (1) if the inversion is present in North America and polymorphic, (2) which environmental conditions modulate the inversion karyotype frequencies, and (3) the relationship between inversion karyotype and adult size.

## Methods

### Study species and field sampling

*Coelopa frigida* belongs to the group of acalyptrate flies and occupies a wide geographic range from Cape Cod (USA) to Greenland on the west coast of North Atlantic Ocean and from Brittany (France) to Svalbard (Norway) on the east coast. Both larvae and adult flies are restricted to decomposing seaweed (wrackbed) for both food and habitat.

We sampled about 90–120 *C. frigida* individuals per population at 20 locations within a 3-week period during September and October 2016. The geographic range of locations spanned over 10° of latitude along the North American Atlantic coast (Fig. 1). At each location, adult flies were collected with nets and preserved individually in ethanol or RNAlater. Environment at each population was described by three categories of variables: local wrackbed seaweed composition, local wrackbed abiotic characteristics, and large-scale climatic/abiotic conditions (Table S3). Wrackbed composition was based on an estimation of the relative proportions of Laminariaceae, Fucaceae, Zoosteraceae, plant debris and other seaweed species; the latter category including various amounts of red, brown and green algae that did not belong to the aforementioned families. Wrackbed abiotic characteristics included an estimation of the surface of the wrackbed and a measure of wrackbed depth, internal temperature and salinity. The three latter variables were estimated by averaging five measurements made at randomly-selected locations of the wrackbed with a salinity multimeter Aquaterr EC350.

**Fig 1:**
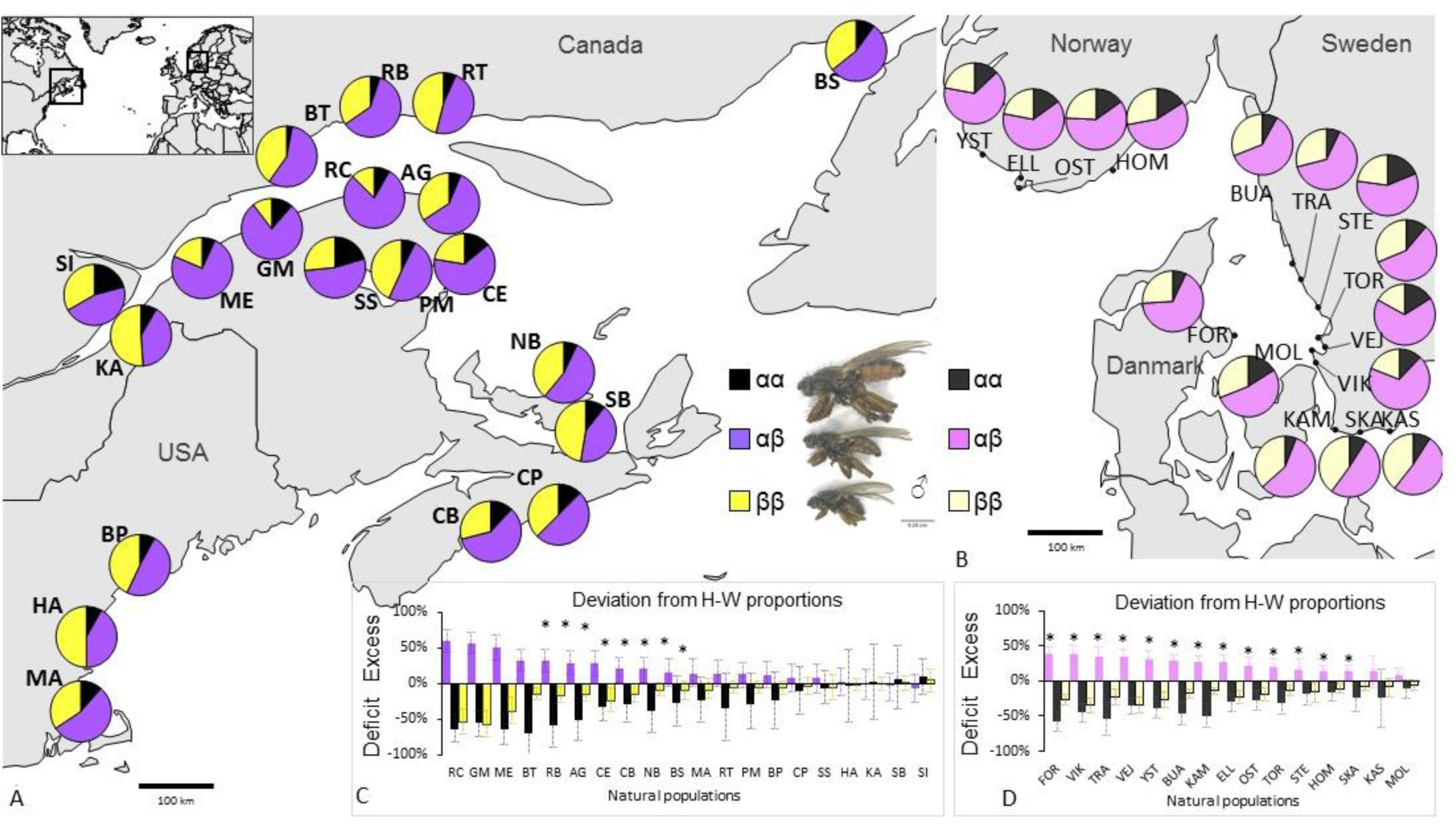
Inversion polymorphism along the North American Atlantic coast and Scandinavian coast. Map of sampled locations and proportions of the three karyotypes at each location in North America (A) and Scandinavia (B). The insert shows the location of the two areas in the North Atlantic area. Deviation from Hardy Weinberg proportions for each karyotype by populations (Calculated as the ratio of observed karyotype frequency over the expected karyotype frequency under HW equilibrium) in North America (C) and Scandinavia (D). Bars represent 95 % confidence intervals and stars denote significant deviation from HW equilibrium (chi^2^ test, p<0.05). All data from Scandinavia are extracted from Day et al 1983, [40].

Large-scale climatic/abiotic conditions were extracted for each location from public databases. These included the annual mean in precipitations and air temperature obtained from the Worldclim database with the R package *Raster* [41,42], the annual mean in sea surface temperature and sea surface salinity obtained from Marspec [43] (except for sites within the St. Lawrence River Estuary for which we used the more accurate local mareograph data of the OGSL, https://ogsl.ca, 2016) and annual mean tidal amplitude. For tidal amplitude, at each location, we extracted hourly water level data from the closest station recorded by NOAA for the US and Fisheries and Ocean for Canada and then calculated the difference between the highest and the lowest water level each day and averaged over the year.

### Fly sex determination, size measurements and inversion genotyping

Adult flies were examined under a binocular magnifier (Zeiss Stemi 2000C) to confirm species identification and to determine sex. For 1967 flies the size was estimated using wing length as a proxy because wings can be easily mounted, flattened for photography, and measured in a standardized way (Fig. S6).

Previous work showed a strong correlation between the chromosome I inversion karyotype (α/β) and two alleles (B/D) of the alcohol dehydrogenase (Adh) allozyme [44]. We used this association to develop an inversion-specific DNA marker and targeted three coding regions within the inversion *(Adh* and two adjacent loci) on which we analysed linkage disequilibrium (LD) and haplotypic variation in American and European samples. LD was calculated as a squared allelic correlation R^2^ [45] between unphased polymorphic sites (including only biallelic site with frequency higher than 5%), tested with chi^2^ test and visualized using the R package *LDheatmap* [46]. Haplotype phasing was inferred using coalescent-based Bayesian methods in DNAsp [47]. Haplotype networks were constructed with median-joining in Network 5.0.0 (http://www.fluxus-engineering.com).

The DNA marker consisted of two single-nucleotide polymorphisms that were associated with the different inversion rearrangements, and these were genotyped with two restriction enzymes (whole procedure detailed in part 1 of Supplementary Materials, Table S1, Fig. S1). It was validated with 44 samples previously karyotyped with the allozyme procedure as described by Edwards *et al* [38] (Table S2) and subsequently used to characterize the karyotype of 1988 wild North American samples of *C. frigida* (89–117 individuals/population, Table S4). For each population, the frequency of α rearrangement and the proportion of each karyotype was calculated in males and females separately, and then estimated for both sexes pooled at a sex-ratio of 1:1.

## Statistical analyses

### Inversion frequencies and Hardy-Weinberg equilibrium (HWE)

Heterogeneity in inversion frequencies and karyotypes frequencies was tested using an analysis of deviance on a generalized linear model (GLM) with binomial logistic transformation, followed by a comparison of contrasts, as well as a pairwise chi^2^ test on proportions between all populations with p-values adjusted following [48]. Within each population, HWE was tested using a chi-square test. Meta-analysis of HWE was tested on this set of p-values using weighted Z-method. Deviation from HWE were calculated for each karyotype as the ratio of observed frequency over the expected frequency. Confidence intervals were drawn by bootstrapping (Table S4).

### Association inversion frequency/environment

Correlation between environmental variables was tested with a Pearson correlation test. For correlated environmental variables within the same category (p<0.05, Pearson R^2^> 0.4), a summary variable was drawn by retaining the first significant PC of a principal component analysis (PCA) on original variables relying on the Kaiser-Guttman and Broken Stick criteria (Fig. S4) [49]. Using the summary variables, VIF (variance inflexion factor) was lower than 2.72, indicating the absence of multicollinearity.

Associations between inversion frequencies and environmental variables were first tested for each variable alone, using a GLM with a logistic link function for binomial data, the response variable being the number of individuals carrying/not carrying the α arrangement and the explanatory variable being an environmental variable, correcting for multiple testing following the Benjamini & Hochberg procedure [48]). Then, the combined effect of the environmental variables was investigated by model selection. For each combination of explanatory variables, two kinds of models were implemented: a GLM, as described earlier, and a beta regression with response variables being the inversion frequency value (bounded between 0 and 1). To identify the best model(s), several indicators were used following [50,51]. First, the models were ranked using AICc values, that is, the small-sample-size corrected version of Akaike information criterion. Second, on the best 25 models, a Jackknife (leave-one-out) procedure was used by repeatedly building the beta-regression model on 19 populations and measuring its predictive fit over the 20 populations. Third, the adjusted R-squared was compared between the most plausible models.

Association between the three karyotypes and environmental predictors was first modelled with a Dirichlet regression. The best Dirichlet model, however, could not predict the relative proportions of the three karyotypes with high accuracy (R^2^ being either small or negative). Comparing the predictive value of the different alternative Dirichlet models showed that each karyotype frequency was best predicted by a different combination of variables (Table S6). Therefore, the karyotype data were translated into binomial proportions (consisting of counts of each category divided by total counts) and analysed separately for each karyotype with binomial GLMs and beta-regression models as described above. As these three models are not independent, they are interpreted accordingly.

Similar analyses were also performed considering as a response variable either the frequencies in males, females or with a sex-ratio as observed in the sampling and led to similar conclusions (Fig. S5, Table S5). We also tested for spatial auto-correlation by building models of redundancy analysis which include environmental predictors and variables describing geographic proximity between populations (Table S7)[52].

For comparison, we re-analysed within the same framework data from Day *et al*. [40] (Fig. 1BD) which examines the cline in the frequency of the inversion in Scandinavia (see part 4 of Supplementary Materials, Fig. S9–10, Table S8–10).

### Relationships between size, inversion type and environment

Size variation in relation to karyotype and sex were analysed with a linear model and post-hoc pairwise t-tests (p-values adjusted following [48]). Residuals from this model are a measure of size variation between individuals, controlled for the combined effect of sex and karyotype and were kept for future analyses (called *“Residuals”)*. Size differences between populations were analysed using a linear model with sex and karyotype as covariates and a post-hoc t-test between each pair of populations on the *Residuals* (p-values adjusted following [48]). The association between size and each environmental factor or inversion frequency was analysed using a linear mixed model with size as the response variable, environmental variables/inversion frequencies as explanatory variables, population as random factor and karyotype/sex as co-variables. Since male mating success may be related to a male size advantage over females (49), we calculated, for each population, the mean size difference between each male karyotypic group and females, and then tested with a linear model whether male-female mean size difference correlated with environmental variables/inversion frequencies.

All analyses were performed in R.3.4.2 [53] using the packages *lme4* [54], *AICcmodavg* [55], *corrplot* [56], *metap* [57], *HardyWeinberg* [58], *lsmeans* [59], *betareg* [60], *vegan* [61], *DirichletReg* [62], *lmertest* [63].

## Results

### Development of a DNA marker of the inversion

The *Adh* gene and the two adjacent coding regions showed a characteristic pattern of low-recombination, consistent with an inversion. First, they were characterized by a very high linkage disequilibrium within and between the three regions over approximately 8 kb with 89% of the SNPs being in significant linkage-disequilibrium (Fig. 2). Second, the three regions showed two distinct haplotype groups, which strongly differed by a total of 41 SNPs and three indels (Fig. 2). In the *Met* regions, 2 haplotypes (out of 62) that elsewhere belonged to the β group shared 11 SNPs characteristics of the α haplotype, suggesting a possible (rare) event of recombination or gene conversion over at least 600bp of the *Met* region. Both α and β haplotype groups included samples from Europe and America. Divergence was stronger between inversion rearrangements (2.4%) than between populations from two continents (0.2%). The haplotype groups and the SNP targeted as a marker showed 100% of concordance with inversion rearrangement karyotypes as determined with the proven allozyme method (4/4 αα, 17/17 αβ, 23/23 ββ, Table S2).

**Fig 2:**
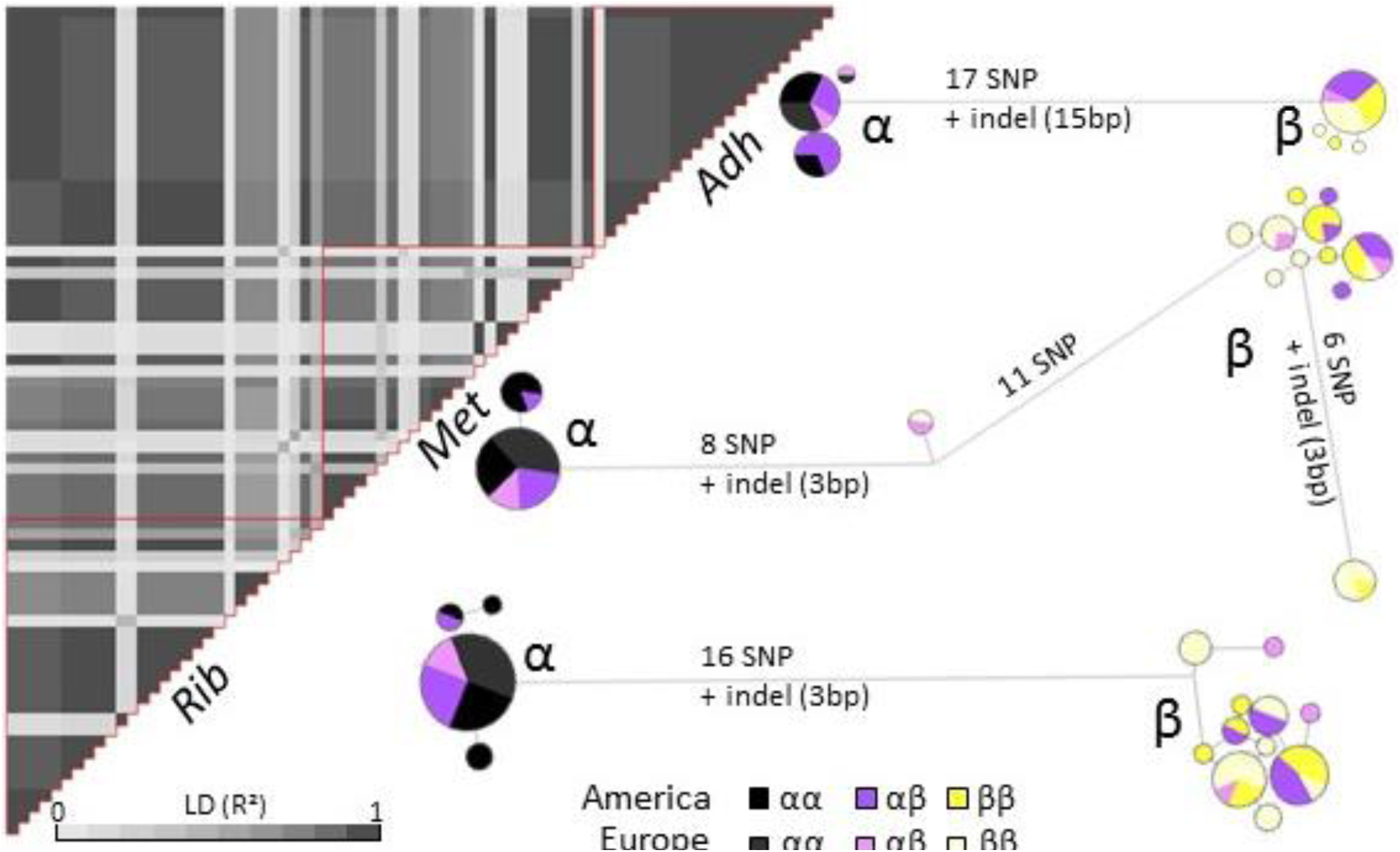
Linkage disequilibrium and haplotype polymorphism. Heatmap representing LD (R2) within and between the three coding regions adjacent to the marker (based on 29 unphased sequences). Haplotypes networks representing, for each coding region, similarity and differences between haplotypes (Adh: 42 samples, 84 haplotypes, Met/Rib: 31 samples, 62 haplotypes). Circles area are proportional to the number of haplotypes with the same sequence. Links are proportional to the number of substitutions. For each locus, two main haplotype groups were found corresponding to the α and the β rearrangements, as labelled. After phasing, heterokaryotypes typically have one haplotype in each group.

### Inversion and karyotype frequencies

All American populations were polymorphic for the inversion and displayed the same global pattern, with α being less frequent than β (α mean frequency=38% [28–51%]) and αα being the rarest karyotype [5–21%] (Fig. 1A). Yet, inversion and karyotype frequencies were significantly heterogeneous between populations (deviance=75(α), 41(αα), 102(αβ), 123(ββ); df=19, p<0.001, Fig. S2–3, Table S4).

Significant deviation from Hardy-Weinberg equilibrium (HWE) was observed among all North American populations (combined probabilities, p<0.001) translating into a mean excess of heterokaryotypes of 20%, due to a mean deficit of 30% of αα homokaryotypes and 15% of ββ (Fig. 1B, Table S4). When considering the 20 populations individually, eight populations showed deviation from HWE with heterozygotes in excess, eight showed a slight excess of heterozygotes (non-significant) whereas four populations were at HWE (Fig. 1B).

### Inversion distribution and environmental variability

North American populations spanned heterogeneous environments, whose variations could be described by two large-scale gradients and heterogeneity in local wrackbed characteristics (Fig. 3A, Fig. S4). The two gradients included a climatic North-South cline, along which co-varied air temperature, sea temperature and precipitations as well as a West-East cline, with lower salinity and higher tidal amplitude in the western part of St Lawrence Estuary. Sampled wrackbeds varied in the seaweed composition, generally dominated by either Laminariaceae or Fucaceae and whose proportions correlated significantly and negatively. Zoosteraceae and plant debris were also present in six out of 20 locations and represented less than 50% (Table S3). The abundance of other seaweed species correlated positively with sea temperature (Fig. 3A). Abiotic characteristics of the wrackbed were split into two independent dimensions, wrackbed surface and a summary variable associating wrackbed depth, with temperature and salinity.

**Fig 3:**
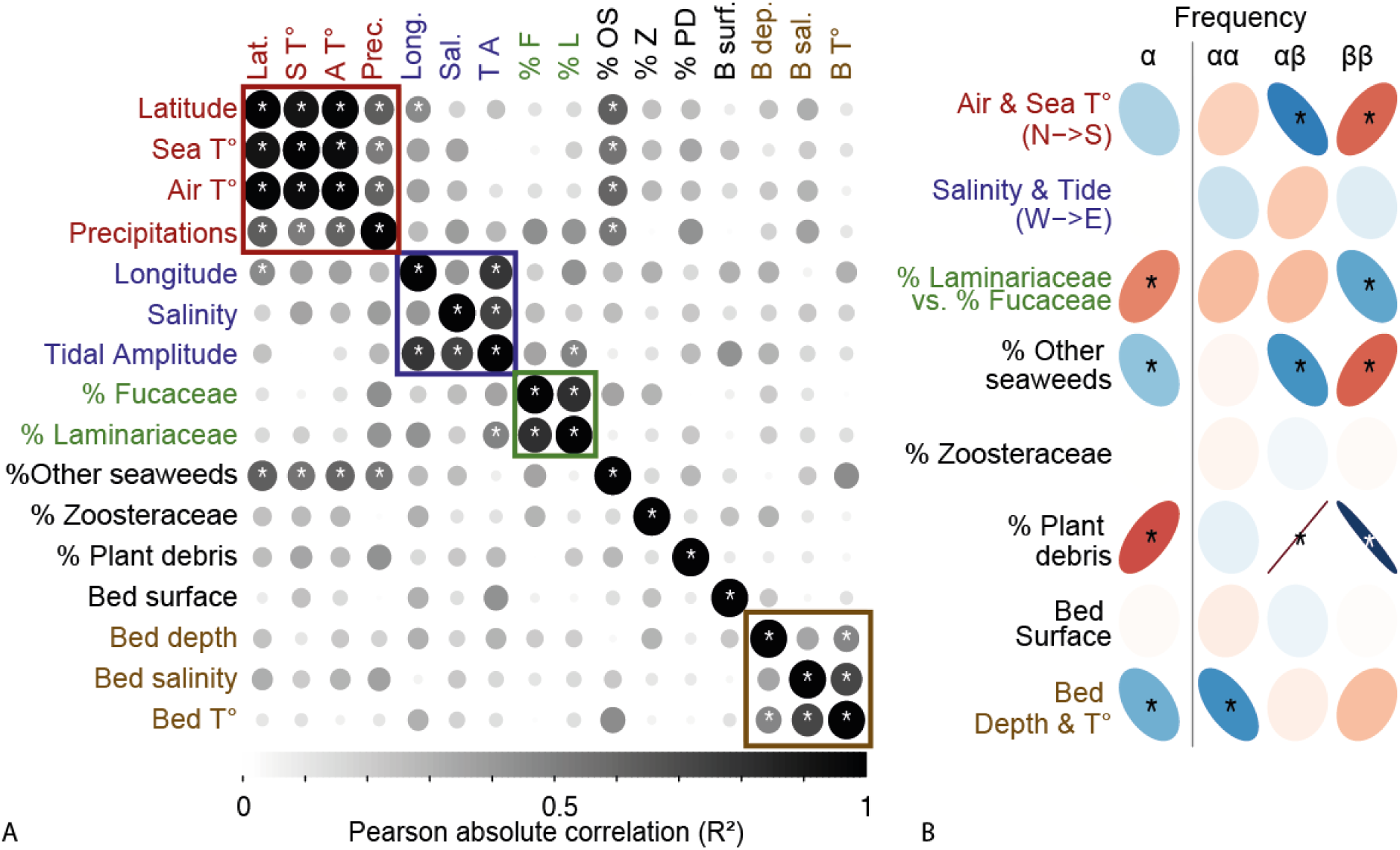
Association between environmental variables and inversion karyotype frequencies. (A) Matrix of Pearson's correlations between environmental variables in North America. Coloured squares delimit groups of variables that were clustered in “summary variables” for subsequent analyses (Fig. S4). (B) Statistical associations between each environmental predictor and the frequency of α rearrangement or the frequency of each karyotype. Strength and direction of the statistical association (GLM) are indicated by the shape of the ellipse and its colour (red: positive, blue: negative). Stars denote significance at 0.05 level, corrected for multiple comparison following [48].

Variation in inversion frequencies were associated with variation in environmental parameters, namely local biotic and abiotic characteristics of the wrackbed, and more marginally, North-South climatic variation. In fact, the best significant predictors of inversion frequency were the composition of the wrackbed and the depth/T° of the wrackbed, which respectively explained 30% and 9% of the variance in the best retained models (Table 1, Table S5). Overall, the α rearrangement was more frequent in shallow and cold wrackbeds, with a high proportion of Laminariaceae or plant debris while the β rearrangement was more frequent in deep and warm wrackbeds dominated by Fucaceae (Fig. 3B). Inversion frequency was also marginally associated with North-South climatic variation, or the correlated presence of other seaweeds, which both explained an additional 3% of variance in alternative best models (Fig. 3B, Table 1, Table S5). The a rearrangement frequency decreased in the South, in warmer areas that contained a high proportion of other seaweed species. This result mirrored the parallel decrease in α frequencies along the Scandinavian North-South thermic cline, in association with warmer air temperature and higher proportion of other seaweeds ([40], re-analysed in Fig. S9–10, Table S8–10).

**Table 1:** Best models explaining the distribution of inversion frequency by a combination of environmental variables

| Model | Beta-regression |  |  | GLM |  |  | R<sup>2</sup><br>adjusted | Jack-knife<br>difference |
| --- | --- | --- | --- | --- | --- | --- | --- | --- |
| | AICc | $\Delta i$ | wAICc | AICc | $\Delta i$ | wAICc | | |
| frequency $\alpha \sim$ %LF + %PD + Bed depth & $T^\circ$ | -48 | 1 | 0.07 | 164 | 0 | 0.10 | 39% | 5% |
| frequency $\alpha \sim$ %LF + %PD | -49 | 0 | 0.10 | 170 | 5 | 0.01 | 30% | 5% |
| frequency $\alpha \sim$ %LF + %PD + Bed depth & $T^\circ$ + %OS | -45 | 4 | 0.01 | 164 | 0 | 0.10 | 42% | 5% |
| frequency $\alpha \sim$ %LF + %PD + Bed depth & $T^\circ$ + Climate | -45 | 4 | 0.01 | 164 | 0 | 0.08 | 42% | 5% |

More detailed analyses to investigate variations in karyotype composition underlying variations in rearrangement frequency showed that each environmental predictor was differentially associated with karyotype proportions (Fig. 3B, Fig. S5, Table 1, Table S5–7). The decrease in α frequency in deeper warmer wrackbeds was linked to a decrease of αα proportions (mostly relative to ββ). The depth/T° of the wrackbed was in fact the best predictor of αα frequency variations (28% of variance). The increase of α frequency with the abundance of plant debris in the wrackbeds was related to higher proportions of αβ karyotypes relative to ββ. Plant debris explained respectively 34% of αβ and 26% of ββ frequency variance. The increase in α frequency with higher proportions of Laminariaceae (vs. Fucaceae) was underlined by higher proportions of αα (and to a lesser extent αβ) relatively to ββ. This variable was a relevant predictor for the three karyotypes (9% of αα variance, 8% of ββ variance, 4% of αβ variance). The association between α frequency and the climatic cline (or the correlated presence of other seaweeds) was mostly due to higher proportions of ββ relatively to αβ karyotypes in the southern part of the cline. Climate and abundance of other seaweeds were slightly redundant but explained respectively 3 and 6% of αβ and ββ variations in alternative models. The positive association between Laminariaceae and αα proportions, as well as the positive association between other seaweeds/warmer air temperature and ββ proportions showed parallelism on the Scandinavian cline (Fig. S9–10, Table S9–10).

Spatial variables were not a significant factor except in one model addressing variation in αα frequency; When controlling for this spatial autocorrelation, the best environmental predictor (depth/T° of the wrackbed) still explained 18% of the variance in αα frequency (Table S7).

Salinity/tidal amplitude was a marginal predictor of the karyotype proportions (9% of αβ variance, 6% of αα variance, 1% of ββ variance,). Yet, it was not associated to variation in α frequency, possibly because the decrease in αβ proportions in the inner estuary was balanced by an increase in both homokaryotypes (Fig. 3B). Wrackbed surface or the proportion of Zoosteraceae were never retained in the best models (Table S5–7).

### Inversion type and size variation

Size was significantly associated with the inversion karyotypes, more strikingly for males than for females (Fig. 4A). For both sexes, the αα karyotype was the largest karyotype and the ββ karyotype the smallest, with heterokaryotypes being intermediates. Size also varied significantly among populations (F_19,1849_=29, p<0.001), with significant differences in 67% (128 out of 190) pairwise comparisons between populations (Fig. S7C). The mean size of each karyotype and sex observed at a given sampling location were significantly correlated (Fig. S7AB), suggesting that local conditions similarly affect size in the three karyotypes and both sexes.

**Fig 4:**
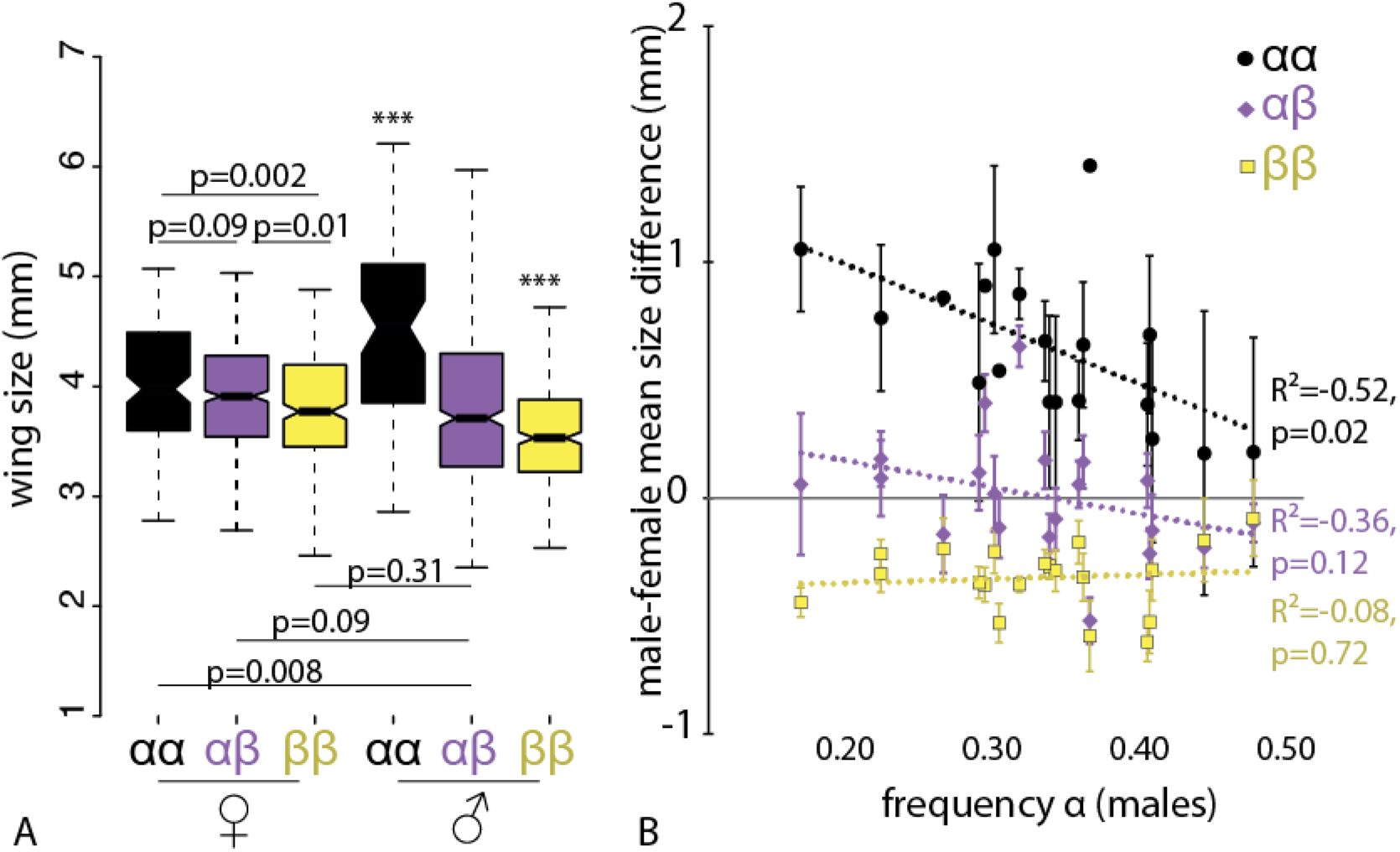
Wing size in relation to karyotype and inversion frequency. (A)Wing size variation by sex and karyotype. Boxes indicate quartile, notches are 95% confidence intervals of the median, whiskers extend to maximal values. *** denotes significant size differences with all other groups (p<0.001). For other comparisons, the p-value of the pairwise t-test is indicated by “p=“. (B)Male-female mean size difference within each population, as a function of α frequency in males. Lines indicate Pearson's correlations (with R^2^ coefficient and p-value of the correlation test).

Size variation was not significantly associated with environment (Fig. S7D). Size variation, controlling for karyotype and sex, was marginally associated with variation in karyotype frequencies with larger flies observed in populations with higher αα frequency (F_1,19_=3.7, p=0.07) and smaller flies found in populations with higher αβ frequency (F_1,19_=3.8, p=0.06, Fig. S7D).

The mean size difference between male and female, a potential indicator of mating success [64], was constant for the smallest ββ karyotype but varied between populations for αα and αβ males, correlating with the α frequency in males (Fig. 4B). Male-female size difference increased at high frequencies of ββ, respectively by a factor of 1 (αα) and 0.7 (αβ), but decreased, by a factor of 3 (αα), at high frequencies of αα (Fig. S8).

## Discussion

Investigating North American natural populations of *C. frigida* revealed the presence of a conserved α/β inversion polymorphism, previously known in European populations of this species [34]. Our results highlighted the importance of local and larger-scale environmental variation in explaining inversion karyotype frequencies, consistent with the prediction that the α/β inversion may contribute to local adaptation [65]. Parallelism in the association between inversion distribution and environment as well as the shared strong haplotype divergence between continents supports the hypothesis that the inversion polymorphism has been subjected to comparable evolutionary processes over an extended range of the species. We discuss hereafter how our results allow new insights into this intercontinental inversion polymorphism and how our data suggests a collective role for several mechanisms of balancing selection.

### A conserved intercontinental inversion polymorphism

As a result of the reduced recombination rate within the inversion, nucleotide sequences within the inversion are generally characterized by high linkage disequilibrium and strong divergence between the different rearrangements [28,66,67]. In *C. frigida*, we identified these characteristics in three adjacent coding regions whose haplotypes were perfectly associated with the *Adh* allozyme marker of the α/β inversion. This shows that recombination is strongly reduced between the two rearrangements and provides the first reliable SNP marker for genotyping the α/β inversion in *C. frigida*. Examining haplotype variations at the three regions further revealed that both haplotypes are found in Europe and America and that haplotype divergence is much stronger than intercontinental variations between populations 5000 km apart. Thus, although further genomic studies are needed to confirm whether this holds true along the inversion, our results point towards a conserved inversion haplotype block throughout the species range. European and American populations also display a similar relationship between inversion karyotype and adult wing size [35] as well as a parallel natural distribution of the inversion [40,44], thus indicating that the α/β inversion in *C. frigida* represents a widespread polymorphism, with similar features conserved throughout the range of the species.

### A role for the inversion in local adaptation to heterogeneous environments

Large climatic gradients or heterogeneous habitats impose spatially-variable selection which favours the evolution of differently-adapted phenotypes, selected for local environmental conditions [1,3]. Inversions are particularly prone to be involved in such local adaptation because they may hold together sets of locally-adapted alleles in the face of gene flow [17,68]. Consistent with these predictions, our results show that *the C. frigida* α/β inversion frequencies co-vary in parallel with the climatic cline and wrackbed composition between both continents (Fig. S9). Although not a hard proof, parallel patterns of genetic or phenotypic variation are considered as strong indirect evidence for local adaptation shaped by natural selection [69].

Along large-scale latitudinal climatic gradients, both *C. frigida* European and American clines exhibit a slight increase of the β arrangement at southern locations where populations experience a higher mean air temperature. This frequency shifts may be a direct effect of increased air temperature, if ββ are less cold-tolerant or if their smaller size/shorter development time is an advantage when warmer temperatures speed up wrackbed decomposition. The latter hypothesis has been used to explain the latitudinal size cline in *Drosophila*, with a smaller size and faster development time being favourable when larval food resources are ephemeral at warmer temperatures [70]. In *C. frigida*, this hypothesis is also supported by the significantly higher frequencies of ββ in warm wrackbeds. The latitudinal cline of frequencies could also be an indirect effect due to the different kinds of seaweeds associated with the southern part of both clines. Admittedly, clinal patterns of variation in allele frequencies could result from isolation by distance without the need to invoke selection. This, however, would unlikely result in similar directionality of the clinal variation across the two continents. Moreover, in Scandinavia, neither population differentiation nor any pattern of isolation-by-distance was observed with SSCP neutral markers over 300km, suggesting that when the habitat is continuous, population structure is weak [71]. This may be surprising considering that the whole life-cycle of *C. frigida* is subjugated to wrackbeds, however occasional mass migratory flight have also been reported which could maintain regular migration between colonies, up to a few hundreds of kilometres [72,73]. Given the relatively continuous habitat observed in North America, little population structure is expected but this needs to be properly tested with neutral markers at a scale appropriate for the cline studied herein (1400km).

Moreover, it is noteworthy that rather than large-scale gradients, the best predictors explaining variation in *C. frigida* inversion frequencies were local wrackbed characteristics, such as the depth, temperature and composition of the wrackbed. Further, the association between those two predictors and inversion frequencies remained when controlling for spatial auto-correlation, suggesting that the environment-karyotype associations are not driven by environmental and genetic similarity between neighbour populations. The influence of the wrackbed composition is of particular interest considering its scale of heterogeneity. The global ratio of Laminariaceae/Fucaceae changes at an order of magnitude along a spatial scale of 100–200 km, a scale at which dispersal is expected [72], and at which local adaptation, related to wrackbed composition, has been observed in Scandinavian populations [65]. Our data suggest that the relative proportions of seaweed species vary with habitat on which each karyotype is preferentially adapted. Both in Europe and America, an increased abundance of Laminariaceae is associated with an increased proportion of αα karyotypes (Fig. S9), a result which is consistent with better survival of αα on Laminariaceae in the laboratory [38]. Mixed wrackbeds or accumulations of plant debris favours heterokaryotypes while wrackbeds of Fucaceae or other seaweeds are associated with increased proportions of ββ. The amount of resources available in each substrate may be one of the factors explaining the different karyotype proportions. In fact, in the laboratory, Laminariaceae sustain a greater viability and a larger size than Fucaceae [38], which suggests it is a richer substrate, facilitating the long larval development and the large size of αα. In the wild populations investigated here, higher αα proportions are also found in populations with large flies (all karyotypes/sexes), and insect size is generally a good indicator of larval growth conditions [74]. Adaptation to different substrates has also been found in the inversion rearrangements in cactophilic *Drosophila*, possibly linked to host-plant specificities in chemical compounds or microbiome fauna [75].

Interestingly, ecological factors identified as good predictors of *C. frigida* karyotype frequencies at a regional scale can also be variable at a finer scale, between neighbouring beaches, and sometimes even within a single beach. Wrackbed depth and temperature, can be patchy; the deposition of plant debris is usually linked to nearby river or storm events. All of these aforementioned factors also vary along a temporal axis. Further work is needed to test whether the association between ecological predictors and karyotypes proportions observed at a regional scale hold true also at a finer scale, *i.e*. within a heterogeneous wrackbed or between neighbouring beaches. If each of the three karyotypes is differentially favoured in each micro-habitat, then they may represent a form of specialisation maintained by micro-spatially varying selection balanced by gene flow. Such kind of balancing selection between micro-niches has been proposed in *Timema cristinae* stick-insects, in which an inversion underlies a green morph and a dark morph, respectively cryptic on leaves or stems of the same host-plant [28]. With a scale of micro-habitat heterogeneity below dispersal distances, inversion structure can be even more important by linking together several adaptive alleles in the face of gene flow, and facilitating the coexistence of different adaptive ecotypes.

### Additional mechanisms of balancing selection contributing to inversion polymorphism

Our results associating inversion karyotype frequencies with latitudinal gradients and seaweed habitats mirrors the well-described clines of inversion frequencies established along eco-climatic gradients in *Anopheles* mosquitoes [21], the mosaic of inversion karyotypes associated with soil moisture in *Mimulus guttatus* [24] and along altitudinal gradients in Kenyan *Apis mellifera* bees [25]. Yet, in those examples, inversion rearrangements are almost fixed at the end of the cline or between habitats, which suggest that selection for each rearrangement is locally very strong. In those cases, spatially-heterogeneous selection, balanced by migration, appears to be the main mechanism determining the geographic distribution and polymorphism level of the inversion [26]. In contrast, clinal variation in *C. frigida* is more modest as all populations are polymorphic and frequencies remain at an intermediate range. Such pattern is not unusual in the literature, with clinal variation of frequencies in *D. melanogaster* spanning between 20–40% [20]; yet it also suggests that additional important factors maintain the polymorphism.

Heterozygote advantage is one the earliest explanation for the persistence of genetic polymorphisms in natural populations [76]. In *C. frigida*, our data and the literature suggest a fitness advantage for heterokaryotypes, which is likely the main mechanism underlying the persistence of this widespread polymorphism across space and time in nature [77]. Indeed, αβ is in excess on both continents and experiments in the laboratory show a higher survival of αβ relative to the two homokaryotypes [36,78]. In the case of an inversion, higher viability of heterokaryotypes can be due to the inversion structure itself. Inversion breakpoints can disrupt important genes, something that can be lethal for one homokaryotype, as shown in the case of the Ruff *Philomachus pugnax* [29]. Reduced recombination over such a large segments of the genome may also prevent the purge of deleterious effects within each rearrangement [30]. For instance, in fire ants *Solenopsis invicta*, one homokaryotype is lethal because of the accumulation of repetitive elements or deleterious mutations [79]. In the case of *C. frigida*, none of the homokaryotypes is lethal but the parallel, repeatedly striking deficit of the homokaryotypes suggests the presence of moderately deleterious effects. This is supported by evidence for genic selection in a series of inter and intra-population crosses [80] and remains to be investigated at the genome level.

Variability in the excess of heterokaryotypes (from 0–60%) indicates that heterozygote advantage may be modulated by local biotic and abiotic conditions. For instance, higher proportions of αβ correlated with particular substrate such as the occurrence of plant debris, or with smaller sizes for all karyotypes/sexes, the latter relationship being probably mediated by density. In the laboratory, smaller size is linked to higher density of larva and density increases overdominance, with a 2.6-fold viability difference between heterokaryotypes and both homokaryotypes at high density but only 1.2-fold at low density [36]. Why heterokaryotypes are better competitors in some environmental contexts is unknown and remains to be investigated.

Heterokaryotype advantage can also result from reproduction [81]. Notably, analyses of wild *C. frigida* female progeny suggest an excess of disassortative mating relatively to inversion karyotype, which may contribute to heterokaryote excess [82]. Polymorphism may also be maintained by opposing viability and sexual advantage. In Soay Sheep *(Ovis aries)* for instance, one allele confers higher reproductive success while the alternative allele increases survival, resulting in increased overall fitness in the heterozygote [83]. This mechanism has also been suggested in *C. frigida* [64]. Because adult size is a major determinant of fertility, larger females lay more eggs, larger males obtain more copulations and both live longer, resulting in αα (and αβ) having a sexual advantage over ββ, particularly in males [39]. Yet, larger size and the associated longer development time in males may not be easily achieved under low-resource conditions, competition or in ephemeral wrackbeds, giving ββ (and αβ) an egg-to-adult viability advantage over αα [36,64]. Such opposing selective pressures may explain the overall heterokaryotype advantage but also the fluctuations of frequencies between low-resource or high-resource substrate. Moreover, in populations with high ββ proportions, αα (and αβ) males are not only rarer, facing less adult competition from large – size males, but they are also larger, possibly because of lower larval competition from similar karyotypes. Thus, α rearrangement could benefit from the “advantage of the rare”, a form of frequency -dependent selection, which is frequently involved in protecting polymorphisms (32). By contrast, in populations with high αα proportions, the observed lower male-female size difference and the numerous same-size competitors are expected to reduce αα males sexual advantage. As such, results from previous studies on European populations along with this study raise the hypothesis that the three karyotypes found in *C. frigida* may represent alternative life history strategies with different relative investment in the trade-off between growth and reproduction and for which balanced polymorphism could be maintained by a form of negative frequency-dependence selection [84] something that could be experimentally tested.

## Conclusion

Our study shows that the αβ inversion polymorphism is conserved between Europe and North America. Significant associations between karyotype frequencies and environmental variables both in North America and Europe provide strong indirect evidence for the role of the α/β inversion in local adaptation. As such, *C. frigida* represents an excellent system to elucidate the multifarious evolutionary mechanisms involved in the maintenance of structural variants. Our data indicates that the three inversion karyotypes are differently favoured by ecological conditions and raises the hypothesis that that may represent three alternative life history strategies, particularly in males. This joins theoretical predictions and accumulating evidence that within-species diversification and specialization can be made possible by the genomic architecture of the inversion itself [14]. Future work in *C. frigida* will focus on population genomics to investigate the contribution of drift and demographic factors, to assess the age of the inversion and to identify which loci are the targets of selection and how linkage increases (or decreases) fitness, depending on the selective process involved. Our analysis combined with the abundant life-history literature on *C. frigida* suggests that several balancing selection mechanisms (e.g. heterosis, genic selection, antagonistic sexual/natural selection, spatially varying and negative frequency-dependent selection) interact to maintain this polymorphism. Further modelling could help to disentangle the relative contributions of these processes in shaping geographic patterns of inversion frequencies and spatial associations. Interestingly, several recent studies also highlighted the combined effects of several balancing selection mechanisms on inversion polymorphism [28,85]. This indicates that the specific architecture of inversions may make them more likely, compared to single-locus polymorphism, to be subjected to multiple and opposing selective factors, and asks under which conditions this results in transient polymorphisms, long-term polymorphisms or speciation.

## Authors’ contribution

CM designed the study, collected field data and samples, did part of the molecular lab work and wing photography, analysed sequences, performed the statistical analyses and drafted the manuscript. CB contributed to molecular lab work. EN & EB contributed to the development of the marker. MW & LB designed and coordinated the study and helped draft the manuscript. All authors contributed in revising the manuscript and gave final approval for publication

## Acknowledgments

We are very grateful to L. Johnson, E. Tamigneaux, D. Malloch for their help during fieldwork and especially to M. Lionard who sampled in Blanc-Sablon. We thank G. Baigle, from the Statistic service (U. Laval), M. Laporte and J. Létourneau for their help with the statistical analysis as well as the Microscopy service at IBIS (U. Laval), S. Bernatchez and B. Labbé for wing photography. We thank three anonymous reviewers for detailed comments that improved the manuscript. This research was supported by a discovery research grant from the Natural Sciences and Engineering Research Council of Canada (NSERC) to L.B, by the Canadian Research Chair in genomics and conservation of aquatic resources held by L. B. and by the Swedish Research Council grant 2012–3996 to M.W. C. M. was supported by a post-doctoral fellowship from the FRQNT and FRQS. EB was supported by a Marie-Curie fellowship.

